# A preliminary analysis of the association between perceived stigma and HIV-related pain

**DOI:** 10.1101/194191

**Authors:** Antonia L Wadley, Tamar Pincus, Michael Evangeli

## Abstract

**Objective**

HIV stigma remains common and has been associated with severity of HIV-related symptoms. Associations between HIV stigma and HIV-related pain, one of the most common symptoms in HIV, have not been investigated however. Data from low back pain populations suggest that stigma associates with worse pain intensity and so we hypothesised that the same would be the case in HIV. In a small pilot study we assessed the association between HIV stigma and pain intensity in PLWH with chronic pain while controlling for depression, a well-established correlate of pain.

**Methods**

Mediation analysis was used to assess the effect of depression on the relationship between stigma and pain intensity in a cross-sectional cohort of 50 PLWH and chronic pain (pain most days of the week for > 3 months) recruited in Johannesburg, South Africa. All participants were assessed using: HIV/AIDS stigma instrument – PLWA (HASI-P), 11-point numerical pain rating scale, and the Beck Depression Inventory II.

**Results**

88% (44/50) of participants reported experiencing some form of HIV stigma (HIV stigma scale score ≥ 1). Worst pain intensity and depressive symptoms individually correlated with total stigma score (Spearman’s r = 0.33, p = 0.02 for both). The mediation analysis highlighted that mediation of the relationship by depression was equivocal (b = −0.002, bootstrapped CI −0.02 to 0.00).

**Conclusions**

Whilst these preliminary data are marginal, they do suggest that associations between HIV stigma and HIV-related pain warrant further investigation. Future study should also include potential mechanisms, which may include mediation through depression.

## Introduction

Stigma in HIV remains prevalent. A survey of over 10,000 HIV-positive individuals in South Africa reported that over one-third perceived themselves as having experienced HIV stigma (1). Stigma against people living with HIV (PLWH) can take different forms including enacted stigma, which refers to acts perceived as stigmatizing such as social exclusion or violence, or internalised stigma, whereby prevalent negative attitudes surrounding HIV are internalised and deemed valid by PLWH (2, 3).

Greater perceived enacted stigma has been associated with increased HIV symptom severity (4). Associations between perceived HIV stigma and the experience of pain have not been investigated, however. Pain is one of the most common symptoms associated with HIV affecting between 50-80% of PLWH (5). The pain frequently is of moderate to severe intensity and occurs concurrently at multiple body sites (5). About half of the pain is chronic (pain on most days for > 3 months) (6, 7).

We recently reported that, in contrast to behaviours seen in many other chronic pain states, patients with chronic HIV-related pain reported not disclosing their pain to friends and family (8). The reason for this behaviour was that revealing their pain status may reveal their HIV status and the participants feared that this would lead to them being stigmatised (8). In individuals with chronic low back pain, stigma in the form of feeling guilty and unbelieved was associated with greater pain intensity and disability (9). Additionally, internalised stigma was associated with pain catastrophizing (a cognitive style comprising frequent pain-related thoughts, rumination and feelings of helplessness) (10), and reduced pain self-efficacy (11). Both increased pain catastrophizing and reduced pain self-efficacy are associated with worse clinical outcomes, including increased pain intensity, in pain populations (10, 12).

These data lead us to hypothesise that HIV stigma is positively associated with pain intensity in PLWH with chronic pain. Moreover, given that HIV stigma is associated with greater depressive symptoms (2, 13) and depression is associated with greater pain intensity (14), the hypothesis here was that greater depressive symptoms would mediate the relationship between HIV stigma and pain intensity.

## Methods

### Design

This was a cross-sectional study of patients visiting an HIV Clinic in Johannesburg, South Africa, between April and June 2015. This clinic is a public, tertiary-care facility providing routine and specialist HIV-related care to PLWH in Johannesburg and surrounding areas. Data collection ran between April and June 2015. Ethical clearance was received from the Human Research Ethics Committee (Medical) of the University of the Witwatersrand (clearance no: M140877).

### Participants, inclusion criteria, recruitment

Participants were informed of the study whilst waiting in the queue to see the doctor and if interested in finding out more, were taken to a private room. Participants willing to be recruited and who fulfilled the inclusion criteria provided written informed consent. Inclusion criteria were: ≥18 years old, confirmed HIV-positive, and the presence of pain most days of the week for at least the last three months. Two interpreters fluent in local languages including isiZulu, isiXhosa and Sotho assisted with consent and study procedures.

Educational attainment was ascertained by asking participants which was the highest grade of schooling they had completed. It was then determined retrospectively whether they had completed primary, secondary or tertiary education. Participants’ employment status was determined by asking if they were currently employed full or part-time, or if they were unemployed. Relevant clinical Information regarding participants’ HIV history including time since diagnosis, time on ARVs, nadir CD4 T-cell count and viral load were retrieved from the participants’ medical files.

### Measures

Worst pain intensity in the last week was measured on an 11-point numerical rating scale anchored at 0: “no pain” and 10: “worst pain imaginable’. This scale has good construct validity in a variety of populations (15, 16) including in African second-language English speakers (17).

Depressive symptoms were measured using the ***Beck Depression Inventory (BDI) II*** (18, 19), which is recommended for use in pain studies (20). The tool has 21 questions, which generate a score from zero to three for each question. The total score is thus 63 with a higher score indicating greater depressive symptoms. Scores between 13 and 19 indicate mild depression and a score ≥ 20 identifies moderate or severe depression (21). The BDI II demonstrated good construct validity and internal consistency (α = 0.90) in an HIV-positive South African population (22). The internal consistency of the BDI II was acceptable in this cohort (α = 0.78).

HIV-related stigma was measured using the ***HIV/AIDS stigma instrument – PLWA (HASI-P)***, which was developed and validated in five sub-Saharan African populations including a South African cohort (23). The tool has 33 questions, split in to three sections. Section one includes 21 questions that assess events that have happened as a result of the participant’s HIV status being disclosed (e,g., “Someone mocked me when I passed by”). The second section contains seven questions related to stigma experienced in healthcare settings (e.g., “I was discharged from the hospital whilst still needing care”). The last section assesses participant’s thoughts and feelings on the disclosure of their HIV status (e.g., “I felt that I brought a lot of trouble to my family”). All statements are rated in terms of how often they have occurred in the last three months: ‘never’, ‘once or twice’, ‘several times’, ‘most of the time’. The sections can be further divided into six subsections: ‘verbal abuse’ (Q4, 9, 10, 11-13, 15, 19), ‘negative self-perception’ (Q29-Q33), ‘healthcare neglect’ (Q22-Q28), ‘social isolation’ (Q7, 8, 16-18), ‘fear of contagion’ (Q1-3, 5, 6, 14), and ‘workplace stigma’ (Q20-21). A mean of the scores for each subsection gives a score of zero to three for each subsection. Internal consistency of the subscales ranged from 0.76 – 0.91, and was 0.94 for the whole scale (23). As the subsections were highly correlated with the total Stigma scale score of which they formed a part (see Supplementary Table), we included the total score in the analyses as a proxy for the subsections. The internal consistency for the total score was found to be excellent in this cohort (α = 0.93).

Participants were asked if they had disclosed their HIV status and if so, whether to family, friends, work colleagues or their pastor. They were also asked, if they hadn’t disclosed to someone, why they had chosen not to.

### Statistical analysis

Normally distributed variables were presented as mean (standard deviation) and non-normally distributed variables as median (range). Spearman’s correlations were used to correlate stigma with both worst pain intensity and BDI scores. The PROCESS tool (24) was used to test the hypothesis that depression mediated the relationship between stigma score and worst pain intensity. Fifty participants were recruited, which was sufficient to detect a medium to large effect size with 0.8 power using the percentile bootstrap method (25). Statistical analyses were carried out using GraphPad Prism 6 (GraphPad, CA), SPSS 2.15 (IBM, NY) and the PROCESS procedure for the mediational analysis. P-values ≤ 0.05 were considered significant.

## Results

The cohort of 50 participants is described in Table 1. They were a middle-aged, predominantly female cohort, and had a high level of unemployment. All participants were on stable antiretroviral therapy. Worst pain experienced in the last week was generally severe (7-10 on NRS): the median pain intensity was 8/10, with 62% (31/50) experiencing pain intensity at 8/10 or greater. The majority had more than one pain site, with head and spine being the most frequent sites of pain (both: 40%, 20/50). Forty-eight percent (48%, 24/50) scored >20 on the BDI-II (moderate or severe depression) and 24% (12/50) scored between 13 and 19 (mild depression).

**Table 1.** Demographic characteristics of the cohort.

| **Variable** | **Value**<br>(n = 50) |
| --- | --- |
| Age (years)<sup>†</sup> | 45 (10) |
| Female <sup>&</sup> | 44 (88) |
| Highest level of education* <sup>&</sup> |  |
| No education | 6 (12) |
| Primary | 16 (32) |
| Secondary | 23 (46) |
| Tertiary | 4 (8) |
| Employment status<sup>&</sup> |  |
| Full time | 21 (42) |
| Part time/piece work | 7 (14) |
| Unemployed | 22 (44) |
| Years since diagnosis<sup>†</sup> | 8 (5) |
| Years on antiretroviral therapy (ART) <sup>†</sup> | 5 (3) |
| Nadir CD4 T-cell count (cells.mm<sup>-3</sup>)<sup>‡</sup> | 152 (2 – 426) |
| Number with undetectable viral load<br>(<40 copies/ml)<sup>†</sup> | 39 (78) |
| Worst pain intensity (0 - 10) <sup>‡</sup> | 8 (4 – 10) |
| Number of pain sites<sup>‡</sup> | 2 (1 – 4) |
| Beck Depression Inventory (0 - 63)<sup>†</sup> | 19 (8) |
| Total stigma score (0 – 99) <sup>‡</sup> | 7 (0 - 64) |

† Mean (SD)
‡ Median (range)
& Count (%)
\* Missing data for one participant

Overall, 88% (44/50) participants reported experiencing some form of HIV stigma (Stigma Scale score ≥ 1) in the preceding three months. Anecdotally, whilst completing the stigma tool, participants frequently reported that the enacted stigma statements weren’t relevant to them because they hadn’t disclosed their HIV status. Indeed, whilst 98% (49/50) of participants had disclosed their status to someone they trusted, 62% (31/50) of patients reported that they had specifically not disclosed their HIV status to their remaining family, friends or co-workers because they feared being stigmatized.

### Univariate analysis

The intensity of worst pain in the last week correlated with total stigma score (Spearmans r = 0.33; p = 0.02). Similarly, greater depressive symptoms (BDI score) correlated with total stigma score (Spearmans r = 0.33; p = 0.02).

### Mediation analysis

Mediation analysis was used to determine whether depression mediated the effect of stigma on worst pain intensity. The direct effect of stigma on worst pain intensity was on the threshold of significance in this model (b = 0.03, CI 0.00 to 0.07, p = 0.05). The indirect effect of stigma on worst pain intensity through depression was tested using a bootstrap estimation (n = 1000). The upper limit of the confidence interval was zero, making the interpretation of whether depression mediated the relationship between total stigma and worst pain intensity equivocal (b = −0.002, bootstrapped CI −0.02 to 0.00).

## Discussion

The hypothesis for the study was that HIV stigma would be associated with pain intensity and that depressive symptoms would mediate the relationship. The results showed that HIV stigma was associated with worst pain intensity but whether this relationship was mediated by depression is unclear.

Perceived HIV stigma was found to be highly prevalent in this cohort (88% of participants), which is far greater than the prevalence of enacted (∼30%) and internalised (35%) stigma reported in a general survey of South Africans living with HIV (1). However, because only patients with pain were recruited, it is not clear if HIV stigma is associated with risk of having pain, but this would be a useful objective of future studies.

The majority of participants had mild to severe depressive symptoms, and while the mean BDI score for this cohort [19 (SD 8)] was greater than that reported by healthy controls [9 (SD 8)], it was similar to another cohort of HIV-positive South Africans [17 (SD 12)] (22, 26). Yet, contrary to our hypothesis, the relationship between stigma and worst pain intensity was not mediated by depression. Future work needs to determine which other psychosocial factors might mediate the relationship between stigma and worst pain intensity. In an Australian chronic pain cohort, pain catastrophizing was associated with internalised stigma (11) and a systematic review of psychological symptoms in people with HIV on treatment, identified higher levels of anxiety than other populations with chronic health issues (27). Stigma causes social withdrawal and exclusion (11) and in an experimental setting, mild to moderate social exclusion led to increased pain sensitivity (28). Thus, catastrophizing, anxiety and social support may be psychosocial factors worth including in future studies of HIV stigma and pain. Additionally, recent work suggested that PLWH and chronic pain may maintain their activity levels despite high pain intensity in order to avoid HIV stigma (8). Thus, physical activity could also be investigated as a mediator of the association between perceived HIV stigma and pain intensity.

There were limitations to the study. This was a cross-sectional study and so it cannot be confirmed that stigma leads to pain. Indeed, the reverse is possible, where worse pain intensity could lead to greater perceived stigma. Longitudinal studies are required to explore the relationship between pain and HIV stigma further. Indeed, data from other longitudinal studies suggest worse physical and mental health (both of which may indicate greater presence and intensity of pain) predicted increases in internalised stigma (29). We recruited patients from an HIV clinic at a tertiary hospital that, in addition to routine care, specialises in dealing with patients failing routine antiretroviral therapy. Thus, some patients included in this study may have had worse mental and physical health than a general population of PLWH and by this logic, may have been at greater risk of HIV stigma. The stigma scale we used, the HASI-P, included the constructs of enacted and internalised stigma (3). Fear of anticipated stigma, the anticipation of future experiences of enacted stigma (3), was highlighted by the qualitative answers suggesting that a proportion of participants had not disclosed their HIV status for fear of enacted stigma. Thus, anticipated stigma would be an important factor to assess in future work. Determining the association between each of these subtypes of HIV stigma and HIV-related pain would be an interesting aim of future work. Another limitation was our small sample size. This study was conducted as a pilot study to test our hypothesis that HIV stigma would be associated with intensity of HIV-related pain. Following the mediation analysis, the confidence intervals generated from the association between stigma and pain intensity identified that the effect might be marginal, which highlights that the association needs to be repeated.

In this first assessment of HIV-related pain and stigma, we found an association between perceived HIV stigma and increased pain intensity. The role of depression in mediating this relationship was not clear. Further study is warranted to repeat these findings and determine the mechanisms by which stigma may influence HIV-related pain, including the role of depression. This information is important to inform psychosocial interventions for HIV-related pain. For example, a recent programme piloted in the US for PLWH, pain and depressive symptoms (30), may be more effective if perceived stigma is also addressed. Furthermore, behavioural interventions for managing pain in HIV (31) may need to take into account that stigma may be a barrier to functional coping with pain.

